# Lionfish (*Pterois volitans*) can see UV light: UV Vision in an Invasive Aquatic Predator

**DOI:** 10.1101/2022.09.09.507242

**Authors:** Elizabeth W. Phillips, Karen E. de Wit, Alexander Kotrschal

## Abstract

UV vision is wide-spread across animals. Many coral reef fish species use the reflection of UV light to communicate with conspecifics, as most aquatic predators are UV-blind. This creates a “secret” communication system for prey fish to signal to others while minimizing their risk of predation. Exploitation of this system by a predator that can see UV light would likely help facilitate prey detection and increase predator efficiency. Recently, indirect evidence has emerged that the Indo-Pacific lionfish (*Pterois volitans*), a highly invasive predator in the Caribbean, might be able to see into the UV spectrum. We propose that UV vision might be an important, and so far overlooked, reason for lionfish’s predatory success. Using an associative learning assay, we investigated lionfish’s ability to discriminate UV from non-UV light and found the first behavioral evidence that lionfish can indeed see UV light. We also measured the UV body reflectance of lionfish and found that lionfish reflect UV light, suggesting that lionfish may communicate with conspecifics via UV vision. Further studies should investigate if the UV vision is more widespread in marine predators than originally thought, as well as continue to investigate the function of UV vision in lionfish and other predators.

## Introduction

Animals can see light in a wide range of spectra. Humans see light in the spectrum of 400-700nm; the ultra-violet (UV) spectrum (100-400nm) constitutes a range that humans and most other mammals are blind to, but a wide range of insects, crustaceans, birds, and fishes can detect (Cortesi et al. 2020; Siebeck and Marshall 2001; Tovée 1995). The function of UV vision in these species has been extensively studied and, in general, falls into five broad categories: navigation, foraging, communication, mate attraction, and circadian rhythms regulation (reviewed in Tovée 1995). Of these functional uses, fishes have been shown to use UV vision mainly during mate attraction, while foraging to better detect plankton, and for short range communication between conspecifics (Flamarique 2013; Losey et al. 1999; Rick and Bakker 2008; Siebeck et al. 2010; Siebeck 2004). However, in marine systems, many predatory marine fish show characteristics in their eye morphology that would prevent the absorbance and detection of UV light (Cummings, Rosenthal, and Ryan 2003; Losey et al. 2003; Lythgoe et al. 1994; Siebeck and Marshall 2000). When investigating the UV transmittance of ocular media in coral reef fish, it was found that around half the 130 predators surveyed showed complete blockage of light below 405nm (Losey et al. 2003; Siebeck and Marshall 2000). Of the predatory species whose ocular media did transmit UV light, most limited UV light transmittance to wavelengths above 360nm (the average peak absorbance of short-wavelength receptors found in fish with UV vision), indicating UV sensitivity, but not necessarily specialized UV vision (Douglas and Mcguigan 1989; Losey et al. 2003; Siebeck and Marshall 2000). Blockage of UV light at the eye can increase predators’ prey detection at long distances, as well as prevent long-term damage to the eye caused by UV light (Carleton et al. 2020; Ivanov et al. 2018; Siebeck 2004; Zigman 1993). Prey species also benefit from predators’ lack of UV vision, as it allows for more elaborate coloration in the UV spectrum and therefore increased opportunity for mate attraction and communication with conspecifics, without risk of increased predation, a trade-off well-documented in the visual spectrum (Burk, 1982; Endler, 1987; Magnhagen, 1991). Importantly, 24 species of predatory fish were identified with a high likelihood for UV vision based on UV transmittance of ocular media (transmitted UV light under 355nm). Most species were from the families Apogonidae (Cardinalfish) and Holocentridae (Squirrelfish and Soldierfish); small nocturnal fish that feed mainly on crustaceans and invertebrates. Most likely, these species never evolved to block UV light due to the lack of UV light during the night, unlike diurnal predators that can be harmed by the high levels of UV light during the day (Losey et al. 2003). However, two families of predators, Synodontidae (Lizardfish) and Scorpaenidae (Scorpionfish), that show a high likelihood for UV vision, eat mainly small fish, and hunt nocturnally *and* in daylight (Cure et al., 2012; Esposito et al., 2009; Morris Jr. & Akins, 2009; Morris Jr., Akins, Barse, Cerino, & Freshwater, 2008; Soares, Barreiros, Sousa, & Santos, 2002). These predators also largely attack prey at close range, making them less likely to be impacted by the decreased visual clarity at long distances caused by UV vision, while also potentially benefitting from the ability to tap into the “secret” UV communication channel used by prey fish. Indeed, a freshwater predator with UV ability, the brown trout (*Salmo trutta*), is more likely to attack three-spined sticklebacks (Gasterosteus aculeatus) that reflect UV light than members of the same species that don’t (Modarressie, Rick, & Bakker, 2013).

Lionfish (*Pterois volitans*) are scorpionfish present in coral reefs and are infamous for their incredible invasive success in the Atlantic and Caribbean. Recently, physiological evidence has emerged revealing the potential for UV vision in lionfish. Corneal electroretinography showed that lionfish corneas are sensitive to light in the spectrum 350-620nm, and that lionfish also possess UV-transmissible lenses with 50% transmittance down to 310nm (Hasenei, Kerstetter, Horodysky, & Brill, 2020). We predict that UV vision may play a role in their invasive ability. The lionfish’s invasion began in 2000 with sightings first recorded in Florida, USA. They rapidly spread up the east coast of the United States and south through the Caribbean towards South and Central America over the next decade (Côté, Green, & Hixon, 2013). Another systematically close, ecologically and behaviorally similar lionfish species (*Pterois miles*) has also recently begun invading the Mediterranean after introduction from the Red Sea through the Suez Canal (Bariche, Torres, & Azzurro, 2013; Dimitriadis et al., 2020; Phillips & Kotrschal, 2021). A thorough understanding of what underlies lionfish’s high invasive success is paramount for management efforts, as lionfish invasions have caused dramatic ecological and economic damage to local areas (Côté et al., 2013; Lesser & Slattery, 2011). Their ability to so successfully invade and expand into new habitats is often attributed to their high reproductive ability, lack of natural predators due to venomous dorsal spines, and their generalist diet (Côté et al., 2013; Galloway & Porter, 2019; Hixon, Green, Albins, Akins, & Morris, 2016). We suggest that their hunting technique may also crucially contribute to their invasive success. Lionfish use a variety of techniques when hunting, all of them requiring close proximity of lionfish to their prey (Hixon et al., 2016). This proximity opens up the potential for lionfish to take advantage of the normally secret short distance UV signaling used by prey fish and may hence improve lionfish hunting ability through better prey detection. Furthermore, according to the naïve prey hypothesis, native prey fish in the territories invaded by lionfish may not have prior experience with predators possessing UV vision, so will be naïve to the increased risk posed by lionfish and are more in danger of being predated on (Cox & Lima, 2006; Sih et al., 2010). This is supported by evidence of higher feeding rates and larger growth rates of lionfish in invasive vs. native ranges (Cure et al., 2012), as well as other studies that show the inability of prey fish in the Caribbean to see the lionfish as a predator (D’Agostino et al., 2020).

While physiological evidence of UV transmittance in ocular media and the presence of UV wavelength absorption in visual receptors suggests lionfish possess UV vision, it is currently unknown whether UV light is further processed in the brain and can be functionally used by this species. Behavioral experiments are one way to test for the use of certain visual spectrums in a species (Kelber, Vorobyev, & Osorio, 2003). For example, in a color association learning task, individuals are presented with two or more colors and taught to associate one of those colors with a food reward. If the animals can learn to discriminate between those colors, then they must be able to perceive them as different. Therefore, such an assay can be used to test if animals of one species can distinguish between the cues within the light spectrum of interest. Lionfish possess learning ability similar to other fish species, including in an associative learning assay using colors in the visual spectrum (de Groot, 2021; Deroy, Hussey, & Macisaac, 2020). Here, we use an associative learning assay to determine if lionfish can differentiate between light containing UV and light without UV as a measure of lionfish’s ability to see into the UV spectrum. We also propose that lionfish may use UV vision to communicate with conspecifics, similar to other coral reef fish species. Previous research has shown UV reflectance in fish can be used for species recognition, mate choice, and to evaluate the threat potential of conspecific intruders (Cummings, Rosenthal, and Ryan 2003; Losey 2003; Siebeck et al. 2010; Siebeck 2004; Smith et al. 2002). Such reflectance in lionfish would be a first indication of intraspecific communication via UV coloration in this species.

## Methods

### Animal Husbandry

The lionfish used in this study were supplied by the Coral and Fish Store (Breda, Netherlands), originally caught within their native range in Indonesia. The fish ranged in size from 3–12cm SL at the time of the experiments and were individually housed in 100-liter tanks equipped with a PVC pipe shelter and an air stone. All tanks were connected by a flow-through filtration system and kept at a temperature of 24°C, pH of 8.0, and salinity of 31ppt. The room housing the lionfish was equipped with overhead fluorescent lights (400-700nm), as well as LED lights above the bottom row of tanks (380-740nm), all set to the same 12:12 light/dark cycle. To minimize the impact of daytime lighting conditions on experimental results, the experiments were conducted only during the dark cycle when no overhead lights were on. Fish were fed once a day, five days a week during the first experiment, and twice a week during the second experiment. They were fed with a varied diet of shrimp, mackerel, and cod, and were fed each day until their stomachs were bulging (sign of satiation in lionfish) or until they stopped showing interest in food when presented. The associative learning experiment took place between August 2021 and October 2021 and data collection of UV body reflectance took place in February 2022. Both experiments were conducted at Wageningen University, Netherlands.

#### Experiment 1: Associative Learning with UV and Non-UV Light as Cues

##### a. Experimental Apparatus

The learning apparatuses used were introduced to housing tanks prior to the start of the experiment and left in the tank for the entire duration. The body of the structure consisted of two rectangular compartments, side-by-side, closed off by three walls and a bottom (Figure 1). The side walls were made from black opaque PVC, while the back wall was made from transparent PVC plate. This side was positioned against the tank wall to allow the observer to score associative learning trials in real-time. The front side of the compartment was left open, allowing fish to enter a compartment and make a choice during learning trials, but was closed off with two doors (one opaque and one transparent) outside of trials.

**Figure 1.**
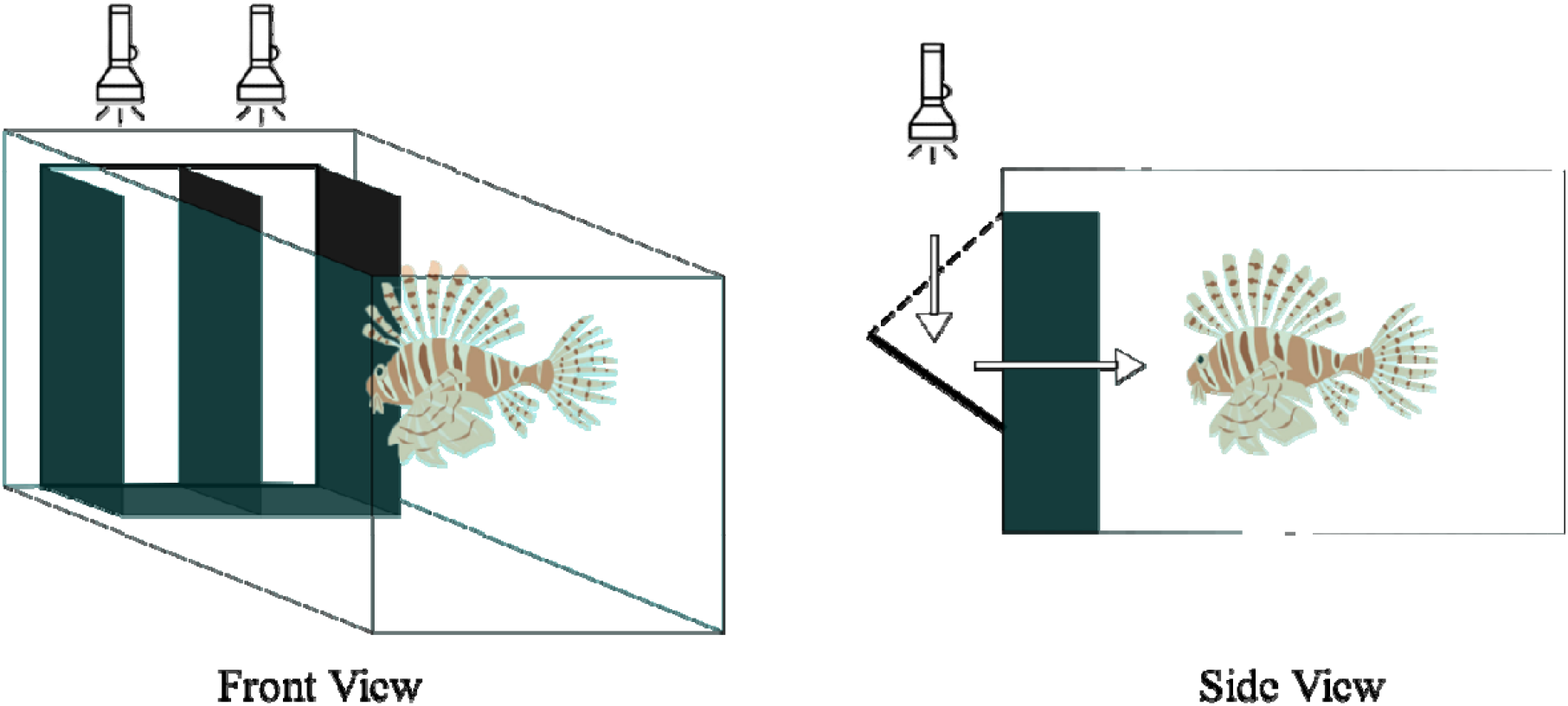
Experimental Apparatus. View of the associative learning apparatus from the fish’s view (left) and the side (right). Black walls mark the opaque walls of the apparatus and the blue walls mark the walls of the tank. Flashlights indicate light source locations from each view, and white arrows indicate the path the light takes when shown to the fish.

During learning trials, light was reflected off of paper and into each compartment to create light cues, as shining light directly into the water disperses it, making spectral differences more difficult to see. Light cues were created using two flashlights (PNQ UV zaklampen) that emit light between 360nm and 395nm. One was covered by a lens (Edmund Optics) that blocks UV light below 390nm, so is referred to as the “non-UV light”, while the other one was left uncovered and is referred to as the “UV light”. When viewed by human eye, the lights shined by the two flashlights seemed identical.

##### b. Pre-training Phase

Prior to the start of experimental trials, lionfish (n=19) were pre-trained to feed from the learning apparatus. During pre-training trials, the doors were lifted one at a time (first the opaque door, then the transparent door five seconds later) and the fish were allowed to inspect the apparatus. If the fish entered either of the two compartments in the apparatus, they were rewarded with food inside the compartment. A fish was considered to have entered a compartment if, when viewing from above, their head up until the base of the pectoral fins was within the walls of the compartment. If after two minutes the fish did not enter the apparatus, a food reward was presented in one of two compartments, determined randomly with the limitation that one side could not be used more than two times in a row. Once food was presented, the trial ended, and the doors were closed. A break of about two minutes was given between subsequent trials. Pre-training continued until the individual swam into either compartment, unprompted, within two minutes for three trials in a row. The eight individuals that did not complete pre-training were not used in the subsequent phase of the experiment. Trials were conducted daily, Monday through Friday. Individuals participated in zero to six trials per day, depending on their hunger level, as trials were stopped each day when the focal fish showed signs of satiation.

##### c. Associative Learning Phase

Once an individual completed pre-training, the next day marked the start of the associative learning phase. Like during pre-training, a trial started when the opaque door of the apparatus was lifted, followed by the transparent door five seconds later. During these trials, UV and non-UV lights were presented as cues for associative learning. Half of the individuals (n=6) were assigned UV light as the rewarded cue, and the other half (n=5) were assigned non-UV light. Each light was randomly assigned to a compartment during each a trial, with the limitation that a cue could not be placed on the same side more than three times in a row for an individual. Individuals were given two minutes to make a choice between the two compartments. If the compartment with the rewarded cue was chosen, this was considered a “correct” choice and food was presented in the compartment as a reward. If the other compartment was chosen, this was considered an “incorrect” choice and individuals were given the chance to correct themselves within the two minutes of the trial. In this case, either food was presented when they entered the rewarded compartment, or at the end of the two minute trial in the rewarded compartment. If no choice was made within those two minutes, food was presented in the compartment with the rewarded cue and the trial outcome marked as “no choice”. Once the food was presented, the trial ended, and the doors were closed. Associative learning trials continued until either the individual reached the learning criteria (seven correct choices in a row), or until the experimental time limit was reached (three months to complete both the pre-training and associative learning phases). The ‘seven in a row’ learning criteria is a common learning criteria in associative learning tasks (Vila Pouca, Mitchell, Lefèvre, Vega-Trejo, & Kotrschal, 2021) as it is significant according to binomial probability.

##### d. Statistical Analysis

To test for learning in the lionfish, we used a one-tailed one-sample Wilcox rank test to determine if the mean success rate of the group was different than what would be expected by chance. Individual lionfish were also tested for learning using a chi-squared test for goodness of fit, comparing the proportion of correct and incorrect responses to chance. Additionally, we also used chi-square tests of independence to determine if lionfish prefer UV or non-UV light by comparing the likelihood of making a correct or incorrect choice in the first trial, first five trials, first ten trials, or overall depending on the light assigned as the rewarded cue. Statistical analyses were conducted in R Studio Version 1.1.463.

#### Experiment 2: UV Body Reflectance

To determine the UV reflectance of lionfish, photos were taken of free-swimming lionfish (n=15) in their tanks in both the visual and UV spectrum. Similar to Kodric-Brown & Johnson (2002), we took photos using filters that either block or absorb light in the UV spectrum to create a colored visual spectrum photo and a black & white UV spectrum photo of each lionfish. For all photos, we used a Nikon D40 camera body with Nikon AF 50mm f/1.8D lens. Visual spectrum photos were taken with light shown directly onto the focal fish from above in an otherwise dark room, and with a UV/IR Cut Filter (Kolari Vision) used on the lens to limit the light captured by the camera to 400-700nm. For the UV spectrum photo, light was provided by two UV flashlights (PNQ UV zaklamp) that emitted light between 360nm and 395nm, and a Ultraviolet Bandpass Transmission Filter (Kolari Vision) was used to limit light captured by the camera to 340-380nm. These UV-spectrum photos were then imported into Adobe Photoshop and converted to black & white images to reveal areas of UV reflectance (white) and areas of no UV reflectance (black). Due to limited lighting during shooting, the brightness of some images was adjusted to better visualize the differences between UV and non-UV reflecting areas.

## Results

### Experiment 1: Associative Learning with UV and non-UV Light as Cues

We found that as a group, lionfish chose the correct cue more often than chance (V = 55, p = 0.003), indicating a learned preference for the rewarded cue. However, only three of the eleven individuals tested reached learning criteria (seven correct choices in a row with no incorrect choices) by the end of the experiment (Figure 2). Of these three, two showed learning at an individual level (Fish 2: x^2^ = 4.5, df = 1, p = 0.034; Fish 4: x^2^ = 7, df = 1, p = 0.008; Table 1).

**Figure 2.**
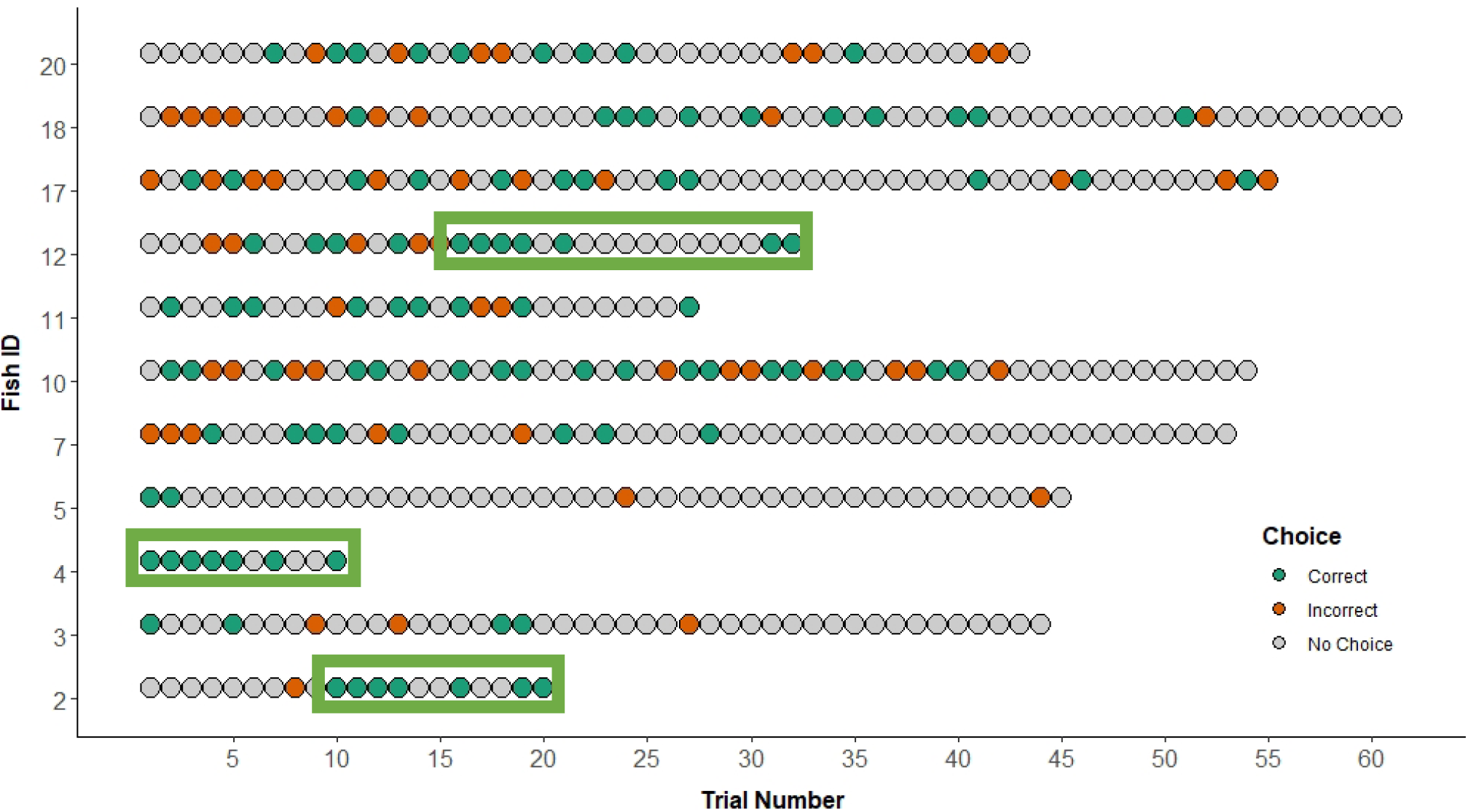
Associative Learning Data. Each circle represents one trial. The color of the circle indicates the choice made by the individual during that trial: green indicates a correct choice, red indicates an incorrect choice, and grey represents when no choice was made. Green boxes are added for individuals that reached learning criteria, and encircle the seven correct trials in a row made by the individual.

**Table 1.** Individual Learning Data. Chi-squared tests for goodness of fit were used to determine individual learning for each fish that participated in the associative learning phase (n = 11). Calculated values for each individual are listed. P-values in bold with an asterisk are significant (p ≤ 0.05).

| Fish ID | Reward Cue | Total Trials | Correct Choices | Incorrect Choices | No Choice | $\chi^2$ statistic | p value |
| --- | --- | --- | --- | --- | --- | --- | --- |
| 2 | UV | 20 | 7 | 1 | 12 | 4.500 | **0.034\*** |
| 3 | UV | 44 | 4 | 3 | 37 | 0.143 | 0.705 |
| 4 | UV | 10 | 7 | 0 | 3 | 7.000 | **0.008\*** |
| 5 | Non-UV | 45 | 2 | 2 | 41 | 0.000 | 1.000 |
| 7 | Non-UV | 53 | 8 | 5 | 40 | 0.692 | 0.405 |
| 10 | UV | 54 | 18 | 12 | 24 | 1.200 | 0.273 |
| 11 | Non-UV | 27 | 9 | 3 | 15 | 3.000 | 0.083 |
| 12 | Non-UV | 32 | 11 | 5 | 16 | 2.250 | 0.134 |
| 17 | UV | 55 | 12 | 11 | 32 | 0.043 | 0.835 |
| 18 | UV | 61 | 11 | 9 | 41 | 0.200 | 0.655 |
| 20 | Non-UV | 43 | 9 | 8 | 26 | 0.059 | 0.808 |

Additionally, we evaluated the effect of reward cue (UV vs. non-UV light) on the likelihood of a fish to make a correct choice during the first trial, first five trials, first ten trials, and overall. We found that in all cases, the assigned reward cue had no effect on the likelihood of fish making the correct choice (first trial: x^2^ = 0, df = 1, p = 1; first five trials: x^2^ = 0, df = 1, p = 0.989; first ten trials: x^2^ = 0.802, df = 1, p = 0.371; overall: x^2^ = 0, df = 1, p = 1).

### Experiment 2: UV Body Reflectance

We found that when photographed in the UV spectrum, all lionfish were visible in photos, indicating they reflect at least some UV light (Figure 3). When comparing UV and visible spectrum photos, we found that the UV-reflective portions of the body match with the white portions of the body in the visible spectrum, while the darker parts do not seem to reflect UV within the spectrum 365-380nm.

**Figure 3.**
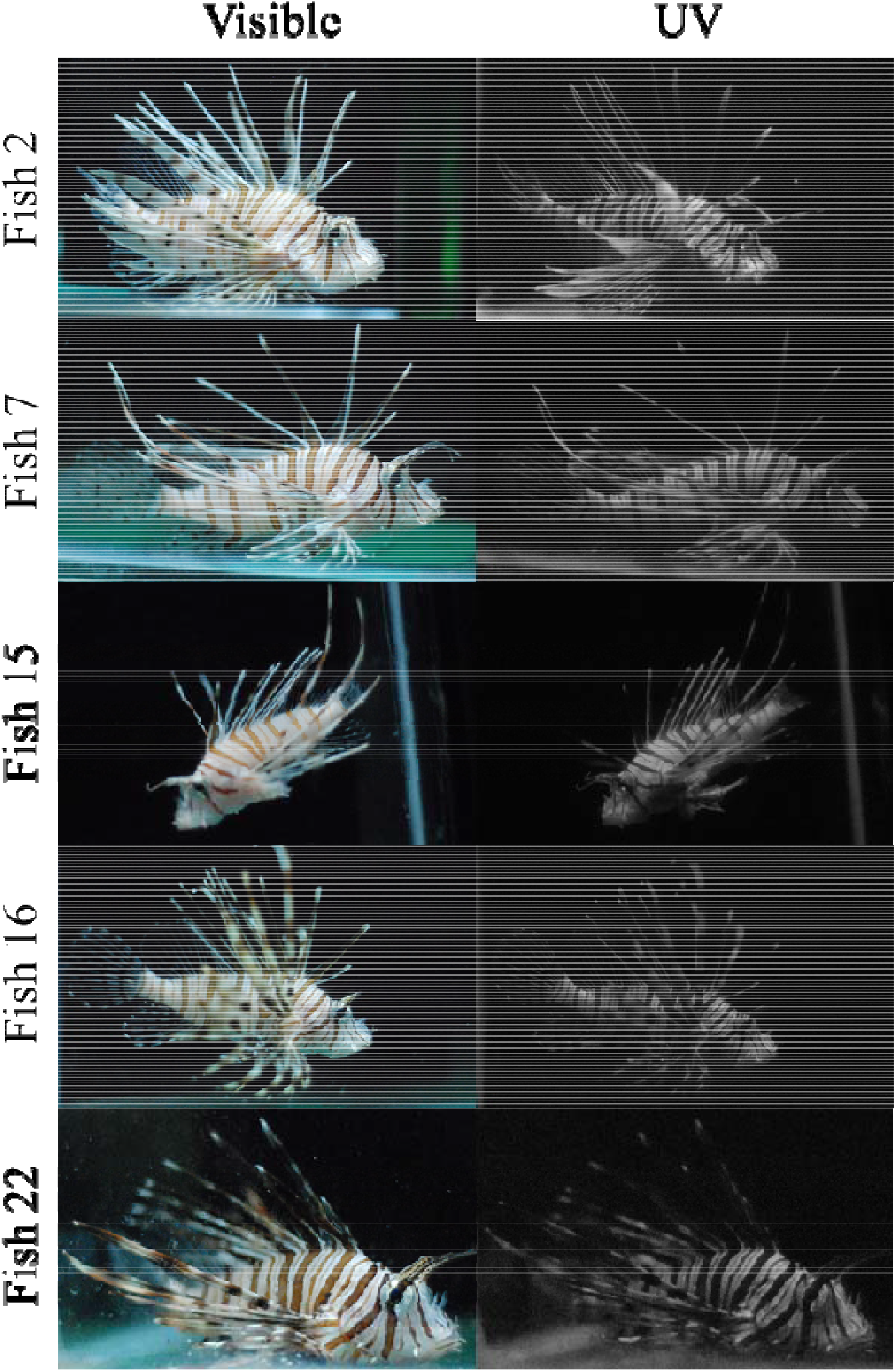
UV Body Reflectance. A subset of lionfish photographed with visual spectrum and UV spectrum photography. Pictures on the left are of lionfish in visual spectrum, and pictures on the right are from the UV spectrum. UV spectrum photos have been converted to black and white for clarity: white area indicate UV reflectance, while black areas indicate no reflectance. Note that the silicone of the fish tank reflects UV light.

## Discussion

Here, we show that lionfish can distinguish between UV and non-UV light using an associative learning task. Overall, lionfish chose the correct cue more often than what would be expected by chance, indicating an ability to distinguish differences between the cues. This is the first behavioral evidence of UV vision in lionfish, and corroborates previous physiological evidence that lionfish can see UV light (Hasenei et al. 2020; Siebeck and Marshall 2001).

We propose that one of the functions of UV vision in lionfish may be to increase hunting efficiency, as many prey fish reflect UV light in coral reefs, a natural habitat of lionfish (Losey et al. 1999; Siebeck and Marshall 2001). One of the ways in which planktivorous prey species use UV vision is for foraging of plankton due to the contrast plankton have with their background in the UV spectrum (Flamarique 2013; Losey et al. 1999). UV light attenuates rapidly in the water column, creating a bright UV-illuminated background against which non-UV reflective plankton are easy to spot. A similar, but opposite effect could explain a potential function of UV vision in lionfish. In contrast to the water column, corals often absorb UV light as a way to mitigate the harmful impacts of UV radiation (Reef, Kaniewska, & Hoegh-Guldberg, 2009). This would provide a darker background on which UV-reflective prey would shine more brightly at close distances. Therefore, lionfish that possess UV vision may have increased hunting success with UV-reflecting prey. Furthermore, prey species in this invasive regions that rely heavily on camouflage as their main predatory defense may not be well-adapted to evade a predator that can see into the UV-spectrum, like the lionfish (Sih et al., 2010). This effect would further increase lionfish hunting ability in invasive regions, and could be a potential cause of the extreme impact lionfish have had on native ecosystems (Cure et al., 2012; D’Agostino et al., 2020; Rojas-Vélez, Tavera, & Acero, 2019). However, as our study only tests the potential of UV vision and not its function for lionfish, additional testing is needed.

One caveat to our hypothesis may be that lionfish typically hunt during the crepuscular period rather than during the day in their native ranges, potentially reducing the effectiveness of UV vision during hunting. However, we believe this does not take away from the potential benefits of UV vision in lionfish for several reasons. In the Caribbean, lionfish have been reported to hunt both diurnally and during twilight hours (Cure et al., 2012; Morris Jr. & Akins, 2009), indicating not only that lionfish sometimes hunt during the day when UV light is most abundant, but also that they may shift their preference for when they hunt based on their environment. According to the naïve prey hypothesis, native prey fish in invaded territories are not well adapted to the predatory strategies of invasive predators and therefore have an increased risk of predation. If prey fish in invasive ranges are less accustomed to predators with UV vision and do not have the skills to avoid predators using this hunting strategy, they can become easier targets for predators like invasive lionfish during the day. We suggest that this may play a role in lionfish’s shift towards diurnal hunting in invasive ranges. In addition, lionfish might still use UV vision to hunt during crepuscular periods, as even though there is less UV light than during the day, there is still some UV light that is reflected by the moon (Robinson, Denevi, Sato, Hapke, & Hawke, 2011). It is unclear how much nocturnal animals use UV light to their advantage, but it remains possible that lionfish utilize UV light during twilight when there is still some amount of light coming directly from the sun, and some reflected off the moon.

We propose that lionfish may use UV vision to also communicate with conspecifics, similar to other coral reef species (Cummings, Rosenthal, and Ryan 2003; Siebeck 2004). To test for the potential of communication with conspecifics via UV vision and reflectance, we used UV photography to take photos of lionfish in the UV spectrum and evaluated their UV reflectance. We found that all the lionfish photographed reflect in the UV spectrum (Figure 3). Notably, all UV-reflecting areas of the lionfish coincided with areas that are white in the visual spectrum, while all UV-absorbing areas were contained within the darker colored regions. This is typical of many UV-reflecting species, as unique patterns that are only visible in the UV spectrum are rare (Losey et al. 1999; Siebeck 2004). Instead, UV-reflectance often enhances the contrast between different areas also visible in the visual spectrum, improving communication through signaling. For lionfish, having this UV reflectance to enhance the already stark contrast in stripes seen in their body coloration within the visual spectrum may increase visibility of their coloration when signaling to conspecifics or during threat displays. It may also increase their crypsis through alternating patterns of UV and non-UV stripes, as is hypothesized to be the purpose of their stripes in the visual spectrum. Alternatively, lionfish UV reflectance may not represent a functional use of their UV vision, but rather be the byproduct of their evolved coloration in the visual spectrum. For example, in one study over half of the investigated species that very likely do not possess UV vision (as light transmittance in their eyes is above 411nm) show reflectance within the UV spectrum (Marshall, Jennings, McFarland, Loew, & Losey, 2003). It is hence clear that additional research is needed to test the functional significance of UV vision, including if lionfish communicate using UV vision and reflectance.

Though our data shows a clear trend of learning at the group level when UV and non-UV cues are used, only two of the eleven individuals tested showed learning at an individual level (Table 1). These low learning rates were most likely caused by the high number of no choices per individual, creating a low number of total trials where choices were made, and may have been due to several reasons. One reason could be that, as this is the attempt to investigate associative learning in lionfish, the learning apparatus may not have been completely optimized for lionfish’s learning ability and feeding ecology, leading to reduced success by individual fish. Indeed, only eleven of the nineteen lionfish that started the experiment made it to the associative learning phase and eight where unable to successfully complete the pre-training. This may indicate that learning to partially enter one of the two compartments in the apparatus to earn a food reward was too difficult for the fish. Alternatively, lionfish might not have been fully motivated to participate in daily trials due to satiation effects, increasing the number of “no choices” in our data set. Another reason the lionfish might have performed more poorly than expected in the associative learning assay could be due to the catching and handling of these fish during catching for the aquarium trade. One of the methods commonly used to catch ornamental fish for the aquarium trade involves using sodium cyanide at sublethal doses to immobilize fish and revive them after capture (Shuman, Hodgson, & Ambrose, 2004). Though sublethal cyanide dosing under laboratory conditions seems to cause no long-term adverse effects in fish, in stressful conditions or at less well-controlled dosages as would be expected during ornamental fish capture, fish show adverse effects of cyanide that are theorized to lead to the 80% mortality rate seen in aquarium fish soon after capture (Hall & Bellwood, 1995; Hanawa, Harris, Graham, Farrell, & Bendell-Young, 1998). Though it was not possible to determine with certainty how the fish in our study were caught, it seems likely that cyanide was involved, and we cannot rule out that cyanide poising may be a reason for the lack of participation seen in our lionfish. One indication for this reasoning may be that smaller fish are typically more susceptible to poison (Reijnders, Aguilar, & Borrell, 2009), and we observed that in our study the relatively smaller lionfish were less likely to succeed during pre-training and ultimately the associative learning assay.

Do the relatively low number of individuals showing clear learning limit the generality of our data? We think not. Studies on primate cognition also use few individuals, as larger samples sizes are difficult to obtain in captivity (Beran, 2008; Bräuer, Call, & Tomasello, 2007; Seed & Tomasello, 2010). In the ‘Alex studies’, even experiments with a single individual Grey parrot (‘Alex’, *Psittacus erithacus*) were used to explore fathom the cognitive abilities of this species (Pepperberg, 2006). As these types of studies investigate the potential reaches of cognition in a species rather than the average level of cognition for a certain population, only a few individuals are necessary to show the capacity for that cognitive ability in a species. In our study, we similarly show that lionfish have the capability to see and discriminate UV light from non-UV light and require only a few individuals to test this hypothesis. Traits such as visual capabilities also do not typically vary much within a species, so from our data it can also be reasonably assumed that if some individuals show the ability to see UV light, most others will as well.

We also noted that the two individuals that showed learning at an individual level were both trained with UV as the rewarded cue. We would expect that if UV light had a functional significance while hunting, lionfish might be more attracted to UV light than non-UV light initially - before an association with a light cue is learned. With our data we could test for such a general preference in the first trial, first five trials, first ten trials, or overall during the associative learning experiment. In all cases, we did not find a significant choice for one over the other, indicating no inherent preference for UV light. However, in our set up, the cues were simply light in an otherwise empty tank. It may be that a setup with more ‘natural’ conditions, e.g. where the reward itself reflects UV light, reveals a preference for UV. Future studies should address this question.

Overall, we show the first behavioral evidence of UV vision in lionfish. This is unique, as most marine predatory fish do not possess UV vision and instead rely on long-distance detection of prey in the visual spectrum. Lionfish, however, capture prey at short-ranges and therefore have the potential to utilize this UV vision to better increase their hunting ability. This may in turn be amplified in invasive ranges, as native prey fish will not have evolved with lionfish to have better defenses to these UV-guided hunting techniques and may therefore be more susceptible to predation by lionfish. More research is needed, however, to determine the function of UV vision in lionfish, whether it be for hunting or communication with conspecifics.

## References

Bariche, M., Torres, M., & Azzurro, E. (2013). The presence of the invasive Lionfish Pterois miles in the Mediterranean Sea. Mediterranean Marine Science, 292–294.

Beran, M. J. (2008). Capuchin monkeys (Cebus apella) succeed in a test of quantity conservation. Animal Cognition, 11(1), 109–116. https://doi.org/10.1007/s10071-007-0094-3

Bräuer, J., Call, J., & Tomasello, M. (2007). Chimpanzees really know what others can see in a competitive situation. Animal Cognition, 10(4), 439–448. https://doi.org/10.1007/s10071-007-0088-1

Burk, T. (1982). Evolutionary Significance of Predation on Sexually Signalling Males. The Florida Entomologist, 65(1), 90–104.

Carleton, K. L., Escobar-Camacho, D., Stieb, S. M., Cortesi, F., & Marshall, N. J. (2020). Seeing the rainbow: Mechanisms underlying spectral sensitivity in teleost fishes. Journal of Experimental Biology, 223(8). https://doi.org/10.1242/jeb.193334

Cortesi, F., Mitchell, L. J., Tettamanti, V., Fogg, L. G., Busserolles, F. De, Cheney, K. L., & Marshall, N. J. (2020). Visual system diversity in coral reef fishes. Seminars in Cell and Developmental Biology, 106, 31–42. https://doi.org/10.1016/j.semcdb.2020.06.007

Côté, I. M., Green, S. J., & Hixon, M. A. (2013). Predatory fish invaders: Insights from Indo-Pacific lionfish in the western Atlantic and Caribbean. Biological Conservation, 164, 50–61. https://doi.org/10.1016/j.biocon.2013.04.014

Cox, J. G., & Lima, S. L. (2006). Naiveté and an aquatic–terrestrial dichotomy in the effects of introduced predators. Trends in Ecology and Evolution, 21(12). https://doi.org/10.1016/j.tree.2006.07.011

Cummings, M. E., Rosenthal, G. G., & Ryan, M. J. (2003). A private ultraviolet channel in visual communication, (November 2002), 897–904. https://doi.org/10.1098/rspb.2003.2334

Cure, K., Benkwitt, C. E., Kindinger, T. L., Pickering, E. A., Pusack, T. J., McIlwain, J. L., & Hixon, M. A. (2012). Comparative behavior of red lionfish Pterois volitans on native Pacific versus invaded Atlantic coral reefs. Marine Ecology Progress Series, 467, 181–192. https://doi.org/10.3354/meps09942

D’Agostino, D., Jimenez, C., Reader, T., Hadjioannou, L., Heyworth, S., Aplikioti, M., … Feary, D. A. (2020). Behavioural traits and feeding ecology Mediterranean lionfish and naiveté native species lionfish predation. Marine Ecology Progress Series, 638, 123–135.

de Groot, C. (2021). Exploring the cognitive landscape of an invasive species: reversal learning in the red lionfish (Pterois volitans). Wageningen University.

Deroy, E. M., Hussey, N. E., & Macisaac, H. J. (2020). Behaviourally-mediated learning ability in an invasive marine fish. Biological Invasions, 0123456789. https://doi.org/10.1007/s10530-020-02329-y

Dimitriadis, C., Galanidi, M., Zenetos, A., Corsini-Foka, M., Giovos, I., Karachle, P. K., … Katsanevakis, S. (2020). Updating the occurrences of Pterois miles in the Mediterranean Sea, with considerations on the thermal boundaries and future range expansion. Mediterranean Marine Science, 62–69.

Douglas, R. H., & Mcguigan, C. M. (1989). The spectral transmission of freshwater teleost ocular media - an interspecific comparison and a guide to potential ultraviolet sensitivity. Vision Research, 29(7), 871–879.

Endler, J. A. (1987). Predation, light intensity and courtship behaviour in Poecilia reticulata (Pisces□: Poeciliidae). Animal Behaviour, 35, 1376–1385.

Esposito, V., Battaglia, P., Castriota, L., Finoia, M. G., Scotti, G., & Andaloro, F. (2009). Diet of Atlantic lizardfish, Synodus saurus (Linnaeus, 1758) (Pisces: Synodontidae) in the central Mediterranean Sea. Scientia Marina, 73(2), 369–376. https://doi.org/10.3989/scimar.2009.73n2369

Flamarique, I. N. (2013). Opsin switch reveals function of the ultraviolet cone in fish foraging. Proceedings of the Royal Society B, 280, 20122490.

Galloway, K. A., & Porter, M. E. (2019). Mechanical properties of the venomous spines of Pterois volitans and morphology among lionfish species. Journal of Experimental Biology, 222(6). https://doi.org/10.1242/jeb.197905

Hall, K. C., & Bellwood, D. R. (1995). Histological effects of cyanide, stress and starvation on the intestinal mucosa of Pomacentrus coelestis, a marine aquarium fish species. Journal of Fish Biology, 47, 438–454.

Hanawa, M., Harris, L., Graham, M., Farrell, A. P., & Bendell-Young, L. I. (1998). Effects of cyanide exposure on Dascyllus aruanus, a tropical marine fish species: lethality, anaesthesia and physiological effects. Aquarium Sciences and Conservation, 2, 21–34.

Hasenei, A., Kerstetter, D. W., Horodysky, A. Z., & Brill, R. W. (2020). Physiological limits to inshore invasion of Indo-Pacific lionfish (Pterois spp.): insights from the functional characteristics of their visual system and hypoxia tolerance. Biological Invasions, 22, 2079–2097. https://doi.org/10.1007/s10530-020-02241-5

Hixon, M. A., Green, S. J., Albins, M. A., Akins, J. L., & Morris, J. A. (2016). Lionfish: A major marine invasion. Marine Ecology Progress Series, 558, 161–165. https://doi.org/10.3354/meps11909

Ivanov, I. V, Mappes, T., Schaupp, P., Lappe, C., & Wahl, S. (2018). Ultraviolet radiation oxidative stress affects eye health, (March), 1–13. https://doi.org/10.1002/jbio.201700377

Kelber, A., Vorobyev, M., & Osorio, D. (2003). Animal colour vision - Behavioural tests and physiological concepts. Biological Reviews of the Cambridge Philosophical Society, 78(1), 81–118. https://doi.org/10.1017/S1464793102005985

Kodric-Brown, A., & Johnson, S. C. (2002). Ultraviolet reflectance patterns of male guppies enhance their attractiveness to females. Animal Behaviour, 63, 391–396. https://doi.org/10.1006/anbe.2001.1917

Lesser, M. P., & Slattery, M. (2011). Phase shift to algal dominated communities at mesophotic depths associated with lionfish (Pterois volitans) invasion on a Bahamian coral reef. Biological Invasions, 13(8), 1855–1868. https://doi.org/10.1007/s10530-011-0005-z

Losey, G. S., Cronin, T. W., Goldsmith, T. H., Hyde, D., Marshall, N. J., & McFarland, W. N. (1999). The UV visual world of fishes: A review. Journal of Fish Biology, 54(5), 921–943. https://doi.org/10.1006/jfbi.1998.0919

Losey, G S, McFarland, W. N., Loew, E. R., Zamzow, J. P., Nelson, P. A., & Marshall, N. J. (2003). Visual Biology of Hawaiian Coral Reef Fishes I. Ocular Transmission and Visual Pigments. Copeia, 3, 433–454.

Losey, George S. (2003). Crypsis and communication functions of UV-visible coloration in two coral reef damselfish, Dascyllus aruanus and D. reticulatus. Animal Behaviour, 66(2), 299–307. https://doi.org/10.1006/anbe.2003.2214

Lythgoe, J. N., Muntz, W. R. A., Partridge, J. C., Shand, J., & Williams, D. M. B. (1994). The ecology of the visual pigments of snappers (Lutjanidae) on the Great Barrier Reef. Journal of Comparative Physiology A, 174(4), 461–467. https://doi.org/10.1007/BF00191712

Magnhagen, C. (1991). Predation Risk as a Cost of Reproduction. Trends in Ecology & Evolution, 6(6).

Marshall, N., Jennings, K., McFarland, W., Loew, E., & Losey, G. (2003). Visual Biology of Hawaiian Coral Reef Fishes. II. Colors of Hawaiian Coral Reef Fish. Copeia, 2003(3), 455–466. https://doi.org/10.1643/01-055

Modarressie, R., Rick, I. P., & Bakker, T. C. M. (2013). Ultraviolet reflection enhances the risk of predation in a vertebrate, 59(2), 151–159.

Morris Jr., J. A., & Akins, J. L. (2009). Feeding ecology of invasive lionfish (Pterois volitans) in the Bahamian archipelago. Environmental Biology of Fishes, 86, 389–398. https://doi.org/10.1007/s10641-009-9538-8

Morris Jr., J. A., Akins, J. L., Barse, A., Cerino, D., & Freshwater, D. W. (2008). Biology and Ecology of the Invasive Lionfishes, Pterois miles and Pterois volitans. Proceedings of the 61st Gulf and Caribbean Fisheries Institute. Retrieved from http://www.e2ccb.org/webpages/gdole/files/LionFish.pdf

Pepperberg, I. M. (2006). Cognitive and communicative abilities of Grey parrots. Applied Animal Behaviour Science, 100(1–2), 77–86. https://doi.org/10.1016/j.applanim.2006.04.005

Phillips, E. W., & Kotrschal, A. (2021). Where are they now? Tracking the Mediterranean lionfish invasion via local dive centers. Journal of Environmental Management, 298, 113354. https://doi.org/10.1016/j.jenvman.2021.113354

Reef, R., Kaniewska, P., & Hoegh-Guldberg, O. (2009). Coral Skeletons Defend against Ultraviolet Radiation, 4(11). https://doi.org/10.1371/journal.pone.0007995

Reijnders, P. J., Aguilar, A., & Borrell, A. (2009). Pollution and Marine Mammals. In Encyclopedia of Marine Mammals (pp. 890–898). https://doi.org/10.1186/1471

Rick, I. P., & Bakker, T. C. M. (2008). UV wavelengths make female three-spined sticklebacks (Gasterosteus aculeatus) more attractive for males. Behavioral Ecology and Sociobiology, 62, 439–445. https://doi.org/10.1007/s00265-007-0471-6

Robinson, M. S., Denevi, B. W., Sato, H., Hapke, B. W., & Hawke, B. R. (2011). LROC WAC Ultraviolet Reflectance of the Moon. In Lunar and Planetary Science Conference (Vol. 6, pp. 15–16).

Rojas-Vélez, S., Tavera, J., & Acero, A. (2019). Unraveling lionfish invasion: Is Pterois volitans truly a morphologically novel predator in the Caribbean? Biological Invasions, 21(6), 1921–1931. https://doi.org/10.1007/s10530-019-01946-6

Seed, A., & Tomasello, M. (2010). Primate cognition. Topics in Cognitive Science, 2(3), 407–419. https://doi.org/10.1111/j.1756-8765.2010.01099.x

Shuman, C. S., Hodgson, G., & Ambrose, R. F. (2004). Managing the marine aquarium trade: is ecocertification the answer? Environmental Conservation, 31(4), 339–348. https://doi.org/10.1017/S03768929040016

Siebeck, U. E., & Marshall, N. J. (2000). Transmission of ocular media in labrid fishes. Philosophical Transactions of the Royal Society B: Biological Sciences, 355(1401), 1257–1261. https://doi.org/10.1098/rstb.2000.0679

Siebeck, Ulrike E., & Marshall, N. J. (2001). Ocular media transmission of coral reef fish - Can coral reef fish see ultraviolet light? Vision Research, 41(2), 133–149. https://doi.org/10.1016/S0042-6989(00)00240-6

Siebeck, Ulrike E., Parker, A. N., Sprenger, D., Mäthger, L. M., & Wallis, G. (2010). A Species of Reef Fish that Uses Ultraviolet Patterns for Covert Face Recognition. Current Biology, 20(5), 407–410. https://doi.org/10.1016/j.cub.2009.12.047

Siebeck, Ulrike E. (2004). Communication in coral reef fish: the role of ultraviolet colour patterns in damselfish territorial behaviour. Animal Behaviour, 68, 273–282. https://doi.org/10.1016/j.anbehav.2003.11.010

Sih, A., Bolnick, D. I., Luttbeg, B., Orrock, J. L., Peacor, S. D., Pintor, L. M., … Vonesh, J. R. (2010). Predatorprey naïveté, antipredator behavior, and the ecology of predator invasions. Oikos, 119(4), 610–621. https://doi.org/10.1111/j.1600-0706.2009.18039.x

Smith, E. J., Partridge, J. C., Parsons, K. N., White, E. M., Cuthill, I. C., Bennett, A. T. D., & Church, S. C. (2002). Ultraviolet vision and mate choice in the guppy (Poecilia reticulata), 13(1), 11–19.

Soares, M. S. C., Barreiros, J. P., Sousa, L., & Santos, R. S. (2002). Agonistic and predatory behaviour of the lizardfish Synodus saurus (Linnaeus, 1758) (Actinopterygii□: Synodontidae) from the Azores. Journal of Ichthyology and Aquatic Biology, 6(2), 53–60.

Tovée, M. J. (1995). Ultra-violet photoreceptors in the animal kingdom: their distribution and function. Trends in Ecology & Evolution, 10(11), 455–460. https://doi.org/10.1016/S0169-5347(00)89179-X

Vila Pouca, C., Mitchell, D. J., Lefèvre, J., Vega-Trejo, R., & Kotrschal, A. (2021). Early predation risk shapes adult learning and cognitive flexibility. Oikos, 130(9), 1477–1486. https://doi.org/10.1111/oik.08481

Zigman, S. (1993). Ocular Light Damage. Photochemistry and Photobiology, 57(6), 1060–1068.

